# Critical spatial separation at the scale of V1 receptive fields determines motion segmentation

**DOI:** 10.64898/2025.12.04.692462

**Authors:** Bikalpa Ghimire, Rohit Bakayat, Abhi Padala, Steven Wiesner, Xin Huang

**Affiliations:** Department of Neuroscience, University of Wisconsin-Madison, Wisconsin 53705, USA

**Keywords:** motion perception, motion integration and segregation, psychometric function, eccentricity, transparent motion, neural mechanism, primary visual cortex

## Abstract

Integrating elements that belong to a single object while segregating overlapping objects is a fundamental challenge for the visual system, exemplified by the phenomenon of motion transparency. While the middle temporal (MT) cortex is central to motion processing, the role of V1 in motion transparency remains controversial. It is still unclear at what spatial scale segmentation for motion transparency occurs. To address these questions, we conducted human psychophysics experiments using locally paired-dot stimuli moving in two directions separated by 90°. Subjects performed a 3AFC task to report whether the visual stimulus had no motion, a single direction, or two distinct directions. We systematically manipulated the path length of the dots, and therefore, the spatial separation between the paired dots, and the retinal eccentricity of the visual stimulus. We found that as the spatial separation between the paired dots increased, subjects’ perception shifted from a single direction to two distinct directions. Critically, we found that the spatial separation required for this perceptual transition increased with the eccentricity and closely matched the known receptive field sizes of V1 neurons at those eccentricities. Direction segmentation occurred only when the spatial separation between motion components exceeded the V1 receptive field size. Conversely, direction integration occurred when the spatial separation was smaller than the V1 receptive field size. Our results demonstrate that the receptive field size of V1 neurons sets the critical spatial scale for direction segmentation and suggest that V1 plays a key role in motion transparency and, more generally, in motion segmentation.

## Introduction

Integrating local elements that belong to the same object while segregating different objects is a universal problem that sensory systems must solve to enable meaningful perception (Braddick, 1993). However, the neural mechanisms underlying these fundamental visual processes remain elusive. Motion transparency refers to the perception of overlapping stimuli moving against each other as segregated surfaces (Clarke, 1977; Marshak and Sekuler, 1979; Andersen, 1989; De Bruyn and Orban, 1993; Braddick, 1997). To generate the perception of motion transparency, the visual system needs to integrate elements with the same motion vector into a single surface while segregating overlapping surfaces from each other. Since the two moving surfaces overlap in the same region of the visual field, motion transparency poses a challenge for the visual system, and at the same time, provides a unique opportunity to investigate the neural mechanisms underlying selective integration and segmentation (Stoner and Albright, 1992; Snowden et al., 1991).

The primate visual system is hierarchically organized (Felleman and Van Essen, 1991). The middle temporal cortex (MT) is a crucial hub for visual motion processing (Born and Bradley, 2005), and receives motion signals from direction-selective neurons in the primary visual cortex (V1) (Movshon and Newsome, 1996; Ungerleider and Desimone, 1986; Maunsell and Van Essen, 1983). It is generally thought that area MT can encode motion transparency based on motion cues (Qian and Andersen, 1995; Andersen, 1997; Braddick and Qian, 2001; Xiao et al., 2014; McDonald et al., 2014; Xiao and Huang, 2015; Huang et al., 2024). However, the role of V1 in motion transparency is controversial (Snowden et al., 1991; Qian and Andersen, 1994). MT neurons have receptive fields (RFs) about ten times larger in size than those of V1 neurons (Gattass and Gross, 1981; Gattass et al., 1981; Albright and Desimone, 1987). A key question is at what spatial scales selective integration and segmentation for motion transparency occurs (Braddick, 1997).

Several psychophysical studies have suggested that, in addition to the motion cues, the spatial arrangement of the motion components can be crucial for motion transparency (Qian et al., 1994a; Curran and Braddick, 2000; Mestre et al., 2001). Qian, Andersen, and Adelson (1994a) introduced locally paired dot stimuli, in which two dots in each pair, moving in opposite directions, were balanced and traveled with a pairing distance of 0.4°. They found that pairing dots locally abolished motion transparency, and human subjects perceived flickering noise. In contrast, when the dots were unpaired, subjects perceived two component directions. These authors suggested that the lack of motion transparency with paired dots moving in opposite directions is due to a local suppression mechanism (Qian et al., 1994a; Qian and Andersen, 1994; Qian et al., 1994b).

Curran and Braddick (2000) further investigated this perceptual phenomenon by using random-dot stimuli moved in two directions separated by 60°, 90°, and 120°. They found that human subjects perceived transparent motion when dots were unpaired, and a single vector average (VA) direction of the motion components when dots were locally paired. Local pairing of dots promotes integration of the motion directions of the paired dots and prevents segmentation of the motion components. The perceptual shift from transparent motion to single VA direction by local pairing of the motion components cannot be explained by a local suppression mechanism (Curran and Braddick, 2000).

Mestre, Masson, and Stone (2001) found that the spatial scale of segmentation based on motion speeds aligns with the receptive field size of V1 neurons. However, the relationship between the size of V1 RFs and the “critical pairing distance” at which perception transitions from integrated single motion direction to segregated component directions has not been established. We hypothesize that when two locally paired dots move in different directions less than 120° apart and travel concurrently within the RFs of V1 neurons, V1 neurons integrate the component directions into the VA direction, resulting the perception of a single motion direction; Conversely, when dots moving in different directions are separated farther than the size of V1 neurons’ RFs, the component directions can be perceptually segregated to generate motion transparency. Our hypothesis predicts that the “critical spatial separation” between dots moving in different component directions, necessary for motion segmentation, should match the size of V1 RFs across retinal eccentricities, and that when the spatial separation between dots is less than the size of V1 RFs, perceptual integration of the component directions should occur.

Here, we conducted human psychophysics experiments to test this hypothesis. We used the paired-dot stimuli moving in two directions separated by 90°, and varied the pairing distance and stimulus eccentricity. We found that as the pairing distance increased, the perception shifted from a single direction to two distinct directions, and the critical spatial separation closely matched the RF size of V1 neurons at different eccentricities. These results have important implications for the neural mechanisms underlying motion transparency and motion segmentation.

## Results

We measured motion perception of human subjects using paired-dot stimuli. The component directions of the paired dots were separated by 90°. Paired random dots moving within a stationary aperture with a diameter of 5°. The dots moved at a constant speed of 5°/s. We manipulated two stimulus parameters. First, we varied the path length (ρ) traversed by each dot, and hence the maximal spatial separation (λ) between the dots in each pair (Fig. 1A, see Methods). The path length was varied from 0.1° to 1.0° in steps of 0.1°, corresponding to maximal spatial separations of 0.071° to 0.707°. Second, we centered the stimulus aperture at retinal eccentricities ranging from 3° to 7° (Fig. 1B). Subjects performed a three-alternative forced-choice (3AFC) task, reporting whether the visual stimulus had no motion, moved in a single direction, or moved in two directions (Fig. 1C).

**Figure 1.**
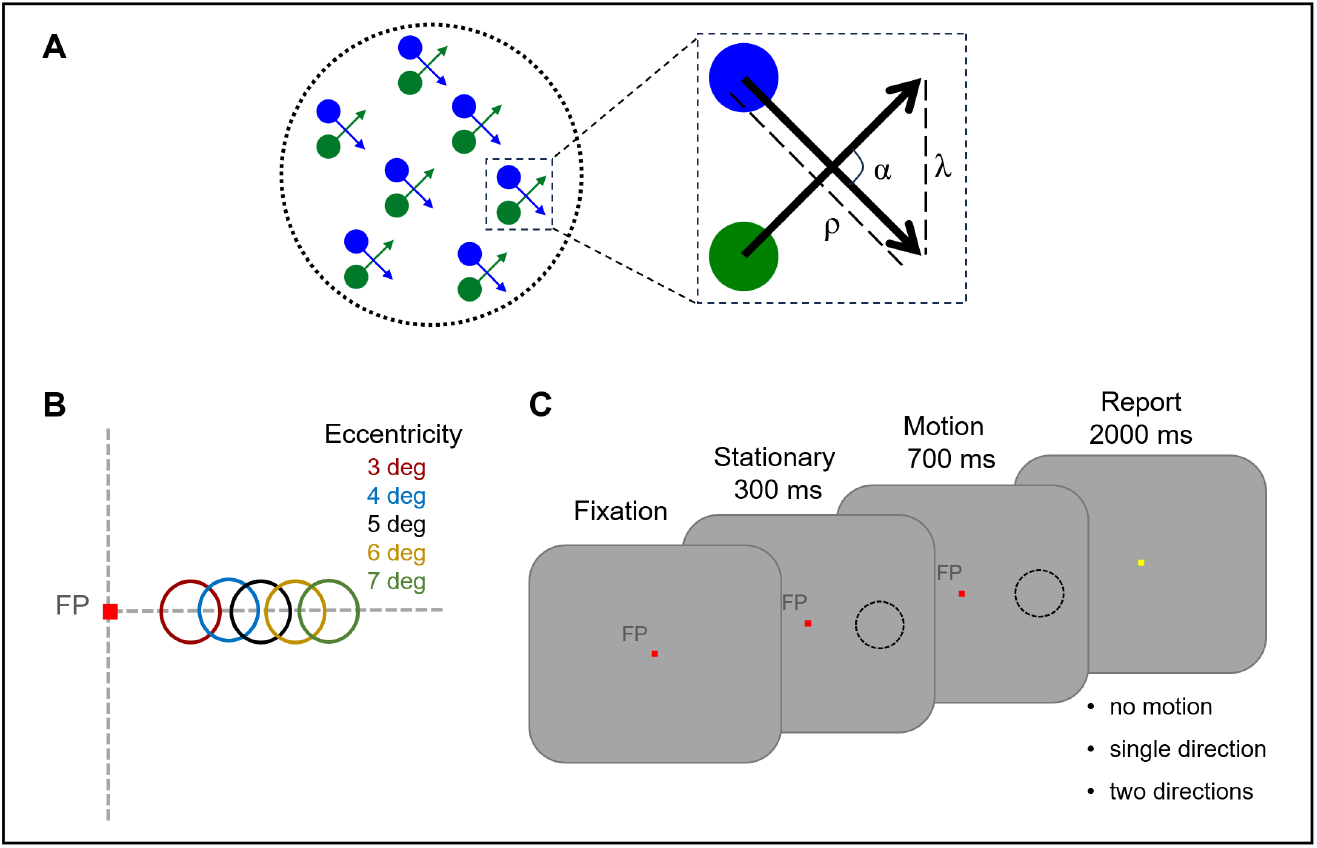
Visual stimuli and experimental paradigm. **A**. A schematic of the paired-dot moving stimulus. Each dot pair consisted of two achromatic dots, shown in blue and green, moving in two directions with an angular separation of α, a path length ρ of the motion trajectory, and a maximum spatial separation λ. The arrow indicates the direction and path length of each dot. **B**. Within each experimental session, the stimulus aperture was centered at five retinal eccentricities of 3°, 4°, 5°, 6°, or 7°. **C**. Experimental paradigm and the 3AFC task. Participants maintained fixation on a central red fixation point (FP) throughout stimulus presentation. After fixation was acquired for 100 ms, the paired-dot stimulus appeared and remained stationary for 300 ms before moving for 700 ms. The vector average (VA) direction of the two component directions was either 0° (rightward) or 180° (leftward), randomly interleaved across trials. Different path lengths of the dots were also randomly interleaved. After stimulus offset, subjects had 2000 ms to report their motion perception by pressing one of three keys to indicate “no motion,” “single direction,” and “two directions.”

### Direction segmentation depends on the spatial separation between motion components

We calculated the proportion of the trials in which subjects reported perceiving two motion directions, and plotted it as a function of the maximal spatial separation of the paired-dot stimulus at each eccentricity (Fig. 2). Taking the result of 3° eccentricity from subject RH as an example, when the maximal spatial separation λ was less than 0.28°, the subject could not perceive two motion directions (Fig. 2A, red curve). As λ increased, the percentage of reporting two directions also increased, rising sharply between 0.35° and 0.49°, reaching nearly 100% at 0.57°. This pattern held for all four subjects, although the slope of the rising phase of the psychometric function and the λ at which the curve asymptoted varied across subjects (Fig. 2B-D). This result indicates that segmentation of two motion directions requires minimal spatial separation between the motion components, and that segmentation performance increases with spatial separation. Also consistent across subjects, the psychometric function shifted to the right at higher retinal eccentricities, indicating that direction segmentation at higher eccentricities requires a larger spatial separation between the motion components (Fig. 2).

**Figure 2.**
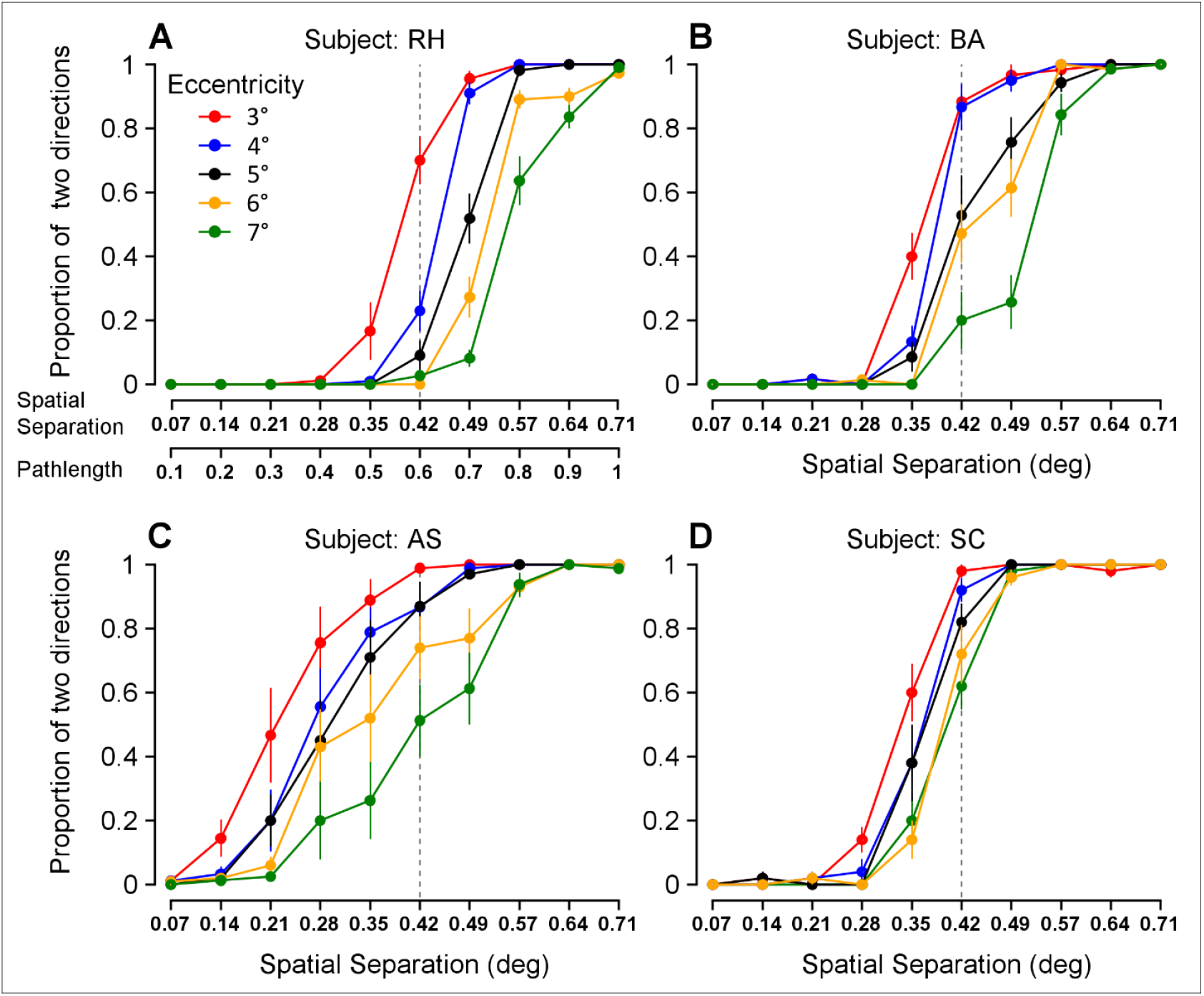
Percentage of perceiving two distinct motion directions as a function of the maximal spatial separation (λ) of the paired-dot stimulus and the effect of retinal eccentricity. **A–D**. Results from four subjects (RH, BA, AS, SC). The abscissa is the maximal spatial separation λ. The corresponding path length (ρ) is also shown in the abscissa of panel A. Color indicates eccentricity of the stimulus. Error bars indicate ± SEM across sessions.

### Critical spatial separation scales with retinal eccentricity

To quantify the relationship between perceptual segmentation and retinal eccentricity, we obtained the critical spatial separation, λ75, defined as the maximal spatial separation between dots in each pair at which subjects reported perceiving two directions in 75% of the trials. To do so, for each subject, we first fit the logistic function to the psychometric function obtained in each experimental session. The logistic function provided an excellent fit to the data in individual sessions (mean *R*^*2*^ = 0.99). We then obtained the λ_75_ value for each session and calculated the mean and variance of λ_75_ across sessions.

We found that the critical spatial separation increased with eccentricity (Fig. 3A). We analyzed this relationship using a linear mixed-effects (LME) model. This approach allowed us to evaluate the fixed effect of eccentricity while accounting for two other sources of variability: differences between individual subjects, modeled as a random effect, and session-to-session fluctuations, modeled as residual error (see Methods). The LME model revealed that the critical spatial separation increased significantly with eccentricity, with a mean slope of 0.036 (95% CI = [0.024, 0.047] and significantly positive, p = 3.652 × 10^-9^) and a mean intercept of 0.251 (95% CI = [0.166, 0.337]) (see Fig. 3A, black regression line). This effect was robust even when accounting for individual differences and session-to-session variation (adjusted R^2^ = 0.671), indicating that eccentricity is the primary predictor of critical spatial separation.

**Figure 3.**
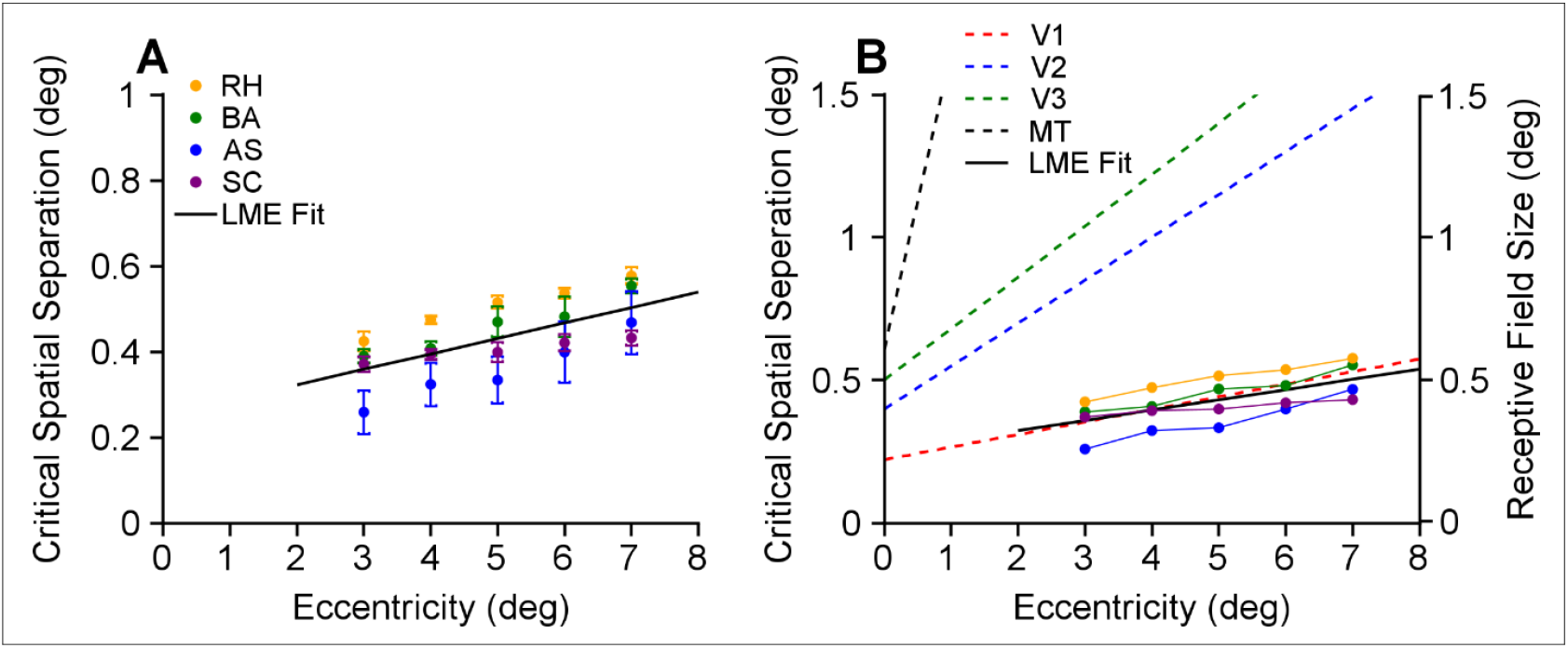
Relationship between critical spatial separation and eccentricity, in relation to the RF sizes of neurons in visual cortices. **A**. Critical spatial separation (λ_75_) as a function of stimulus eccentricity. Data from four subjects, AS, BA, RH, and SC. The symbols represent the mean λ_75_ averaged across sessions for each subject. Error bars represent symmetric 95% confidence intervals (CI) derived from the SD of the bootstrap distribution (1,000 resamples). The solid black line is the LME model fit, y = 0.036x + 0.251 (R^2^ = 0.671). **B**. Relationship between critical spatial separation and RF sizes of neurons at different eccentricities. The λ_75_ values were replotted from panel A, together with the LME model fit (black line). The dashed lines indicate previously published RF sizes of neurons in V1, V2, V3, and MT, as a function of eccentricity.

To confirm that the effect of eccentricity was robust despite this individual variability, we performed two additional analyses. First, one-way ANOVA confirmed a significant effect of eccentricity for every subject (see F-statistic and p-value in Table 1), indicating that critical spatial separation differed significantly across eccentricity. Second, for each subject, linear regression revealed that critical spatial separation and eccentricity had a significantly positive relationship, with eccentricity explaining a substantial proportion of the variance in the data (see p-values and R^2^ in Table 1).

**Table 1.** Testing the effect of eccentricity on critical spatial separation. Summary statistics of linear regression and ANOVA for each subject

| Subjects | Linear Slope | Linear regression $R^2$ | Correlation coefficient (p-value) | ANOVA F statistics | ANOVA p-value |
| --- | --- | --- | --- | --- | --- |
| RH | 0.037 | 0.787 | 0.89 ($p = 4.31 \times 10^{-18}$) | $F(4,46) = 46.07$ | $1.58 \times 10^{-15}$ |
| BA | 0.040 | 0.695 | 0.83 ($p = 1.95 \times 10^{-8}$) | $F(4,24) = 16.16$ | $1.52 \times 10^{-6}$ |
| AS | 0.049 | 0.343 | 0.59 ($p = 1.89 \times 10^{-5}$) | $F(4,41) = 5.76$ | $9.02 \times 10^{-4}$ |
| SC | 0.015 | 0.484 | 0.70 ($p = 1.12 \times 10^{-4}$) | $F(4,20) = 4.93$ | 0.0062 |

### Critical spatial separation aligns with the RF size of V1 neurons

Along the visual hierarchy, areas V1, V2, V3, and MT all have direction-selective neurons and contribute to motion processing (Hubel and Wiesel, 1968; Gegenfurtner et al., 1996; Felleman and Van Essen, 1991; Zeki, 1974). The RF sizes of neurons in these areas progressively increase. We determined which visual area’s RF size corresponds to the scale of spatial separation required for motion segmentation. We compared the critical spatial separation as a function of eccentricity with the known RF sizes of neurons in these areas. Figure 3B shows the RF sizes of V1, V2, V3, and MT neurons measured in previous neurophysiological studies of macaque monkeys (Dow et al., 1981; Gattass et al., 1981; Gattass et al., 1988; Albright and Desimone, 1987). The critical spatial separation as a function of eccentricity, plotted in Figure 3A (slope = 0.036, intercept = 0.251), closely matched the RF sizes of V1 neurons, according to neurophysiological studies (slope = 0.044, intercept = 0.22, Dow et al., 1981). In contrast, the slopes of RF sizes as a function of eccentricity for V2, V3, and MT are 4.17, 5.00, and 28.89 times the critical spatial separation slope measured here, respectively (Dow et al., 1981; Gattass et al., 1981; Gattass et al., 1988; Albright and Desimone, 1987).

### Perceptual motion integration occurs at spatial separations smaller than V1 RFs

The preceding analyses focused on the spatial separations at which subjects perceived segmented motion directions. Curran and Braddick (2000) showed that when the angular separation between two component directions was 60°-120°, human subjects perceived the integrated VA direction for paired-dot stimuli with a path length of 0.2° for at least one motion component. Here, we characterized the range of path lengths and spatial separations within which subjects perceived an integrated single direction, at different retinal eccentricities.

The 3AFC behavioral task allowed us to distinguish between trials in which subjects did not perceive motion and those in which they did. When the path length and corresponding motion duration of the dots were very short, subjects did not perceive motion (Fig. 4). As the path length increased, subjects began to see a single integrated motion direction; The percentage of trials in which subjects perceived a single direction reached a peak, then declined (Fig. 5A-D). We measured the spatial separations at the half-height of the peak percentage during the declining phase, and referred to them as the “spatial separation limit for integration” (SSLI). Only when the spatial separation was less than SSLI and above the threshold for motion detection, did subjects perceive a single motion direction. We found that SSLI increased with the eccentricity and was aligned with the RF sizes of V1 neurons at the corresponding eccentricity (Fig. 5E).

**Figure 4.**
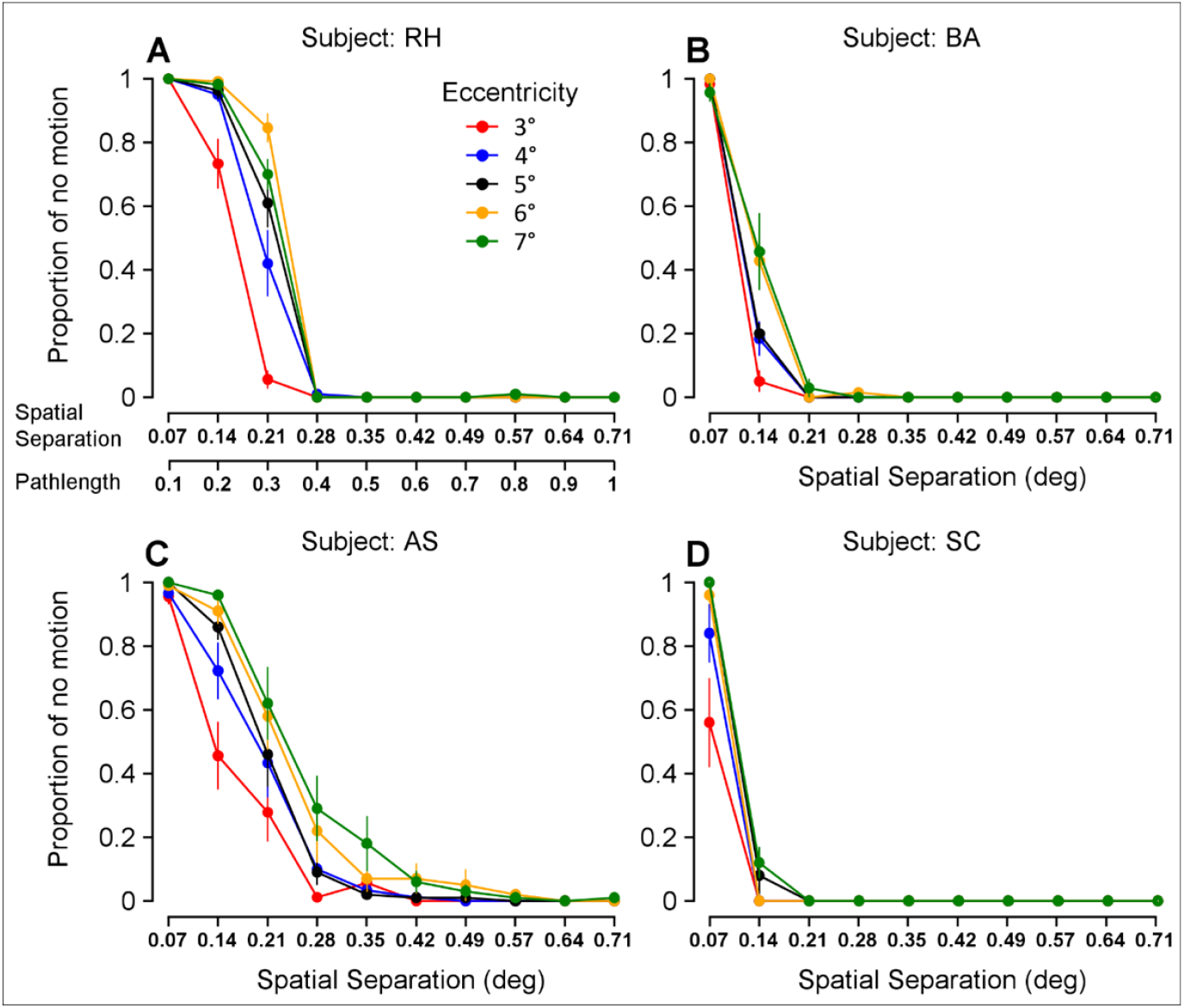
Percentage of reporting no motion as a function of the spatial separation of the paired-dot stimulus and the effect of retinal eccentricity. **A–D**. Results from four subjects (RH, BA, AS, SC). The abscissa is the maximal spatial separation λ. The corresponding path length (ρ) is also shown in the abscissa of panel A. Color indicates eccentricity of the stimulus. Error bars indicate ± SEM across sessions.

**Figure 5.**
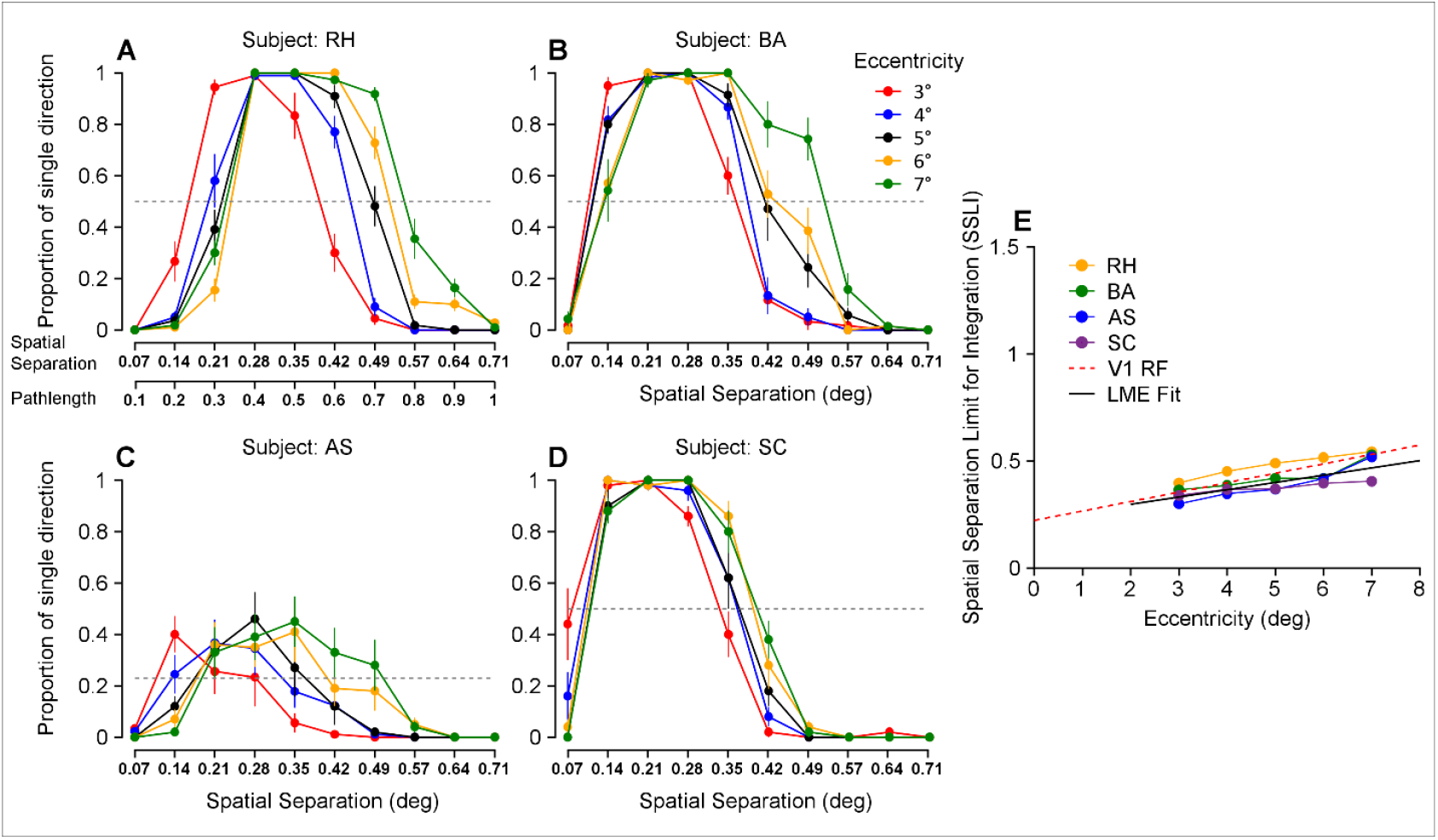
Percentage of perceiving a single motion direction as a function of the spatial separation of the paired-dot stimulus and the effect of retinal eccentricity. Results from four subjects (RH, BA, AS, SC). **A–D**. The abscissa is the maximal spatial separation λ. The corresponding path length (ρ) is also shown in the abscissa of panel A. Color indicates eccentricity of the stimulus. Error bars indicate ± SEM across sessions. **E**. Spatial separation limit for integration (SSLI) as a function of eccentricity. Colors indicate individual subjects. The solid black line is the LME model fit, y = 0.034x + 0.229 (R^2^ = 0.683). The red dashed line shows previously published RF sizes of neurons in V1 based on macaque neurophysiology (Dow et al., 1981).

The LME model revealed that the SSLI increased significantly with eccentricity, with a mean slope of 0.034 (95% CI = [0.023, 0.047] and significantly positive, p = 3.980 × 10^-8^) and a mean intercept of 0.229 (95% CI = [0.144, 0.315]) (see Fig. 5E, black regression line). This effect was robust even when accounting for individual differences and session-to-session variation (adjusted R^2^ = 0.683), indicating that eccentricity is the primary predictor of SSLI. This trend was consistent across all four subjects, despite some individual differences (Fig. 5). Together, these results suggest that perceptual integration of two component directions into a single direction occurs, when two paired dots have a spatial separation less than the RF size of V1 neurons.

## Discussion

Using paired-dot stimuli moving in two directions separated by 90°, we have made several new findings regarding perceptual segmentation and integration of motion directions. First, we showed that, as the pairing distance and, hence, the maximal spatial separation between dots increased, motion perception changed from a single direction to two distinct directions. Second, we found that the critical maximal spatial separation required for direction segmentation increased with stimulus eccentricity. Importantly, the critical spatial separation was well aligned with the RF size of V1 neurons at the corresponding eccentricity, and substantially smaller than RF sizes in extrastriate cortices. Only when the spatial separation between motion components was larger than the RF sizes of V1 neurons, perceptual motion segmentation occurred. Conversely, only when the spatial separation between the paired dots was smaller than V1 RFs, perceptual integration of the component directions into a single direction occurred.

### Relationship with previous studies

Previous studies have not varied the pairing distance of the dots (i.e., the path length each dot travelled) across different eccentricities to capture the transition from an integrated single direction to segregated component directions. In the main experiment of Qian et al. (1994), they used a path length of 0.4°. In the study by Curran and Braddick (2000), they used a path length of 0.2° when the two paired dots moved at the same speed; when the dots moved at different speeds, they fixed the path length of the slower dot to 0.2°. Our study is the first to systematically examine the effect of path length of the dots (hence the spatial separation between motion components) and retinal eccentricity on the integration and segmentation of motion directions.

Mestre et al. (2001) investigated the spatial scale of motion segmentation based on speed cues. They used dot stimuli moving in the same direction at two different speeds and measured the perceptual threshold for speed segmentation. They found that pairing dots at small distances blocked speed segmentation. The critical pairing distance that gave rise to a clear difference in thresholds between paired and unpaired dots at different eccentricities aligned with the RF sizes of V1 neurons. The results of our study on direction segmentation are consistent with those of Mestre et al. on speed segmentation (2001). Together, these findings revealed that, in general, spatial separation at the scale of V1 RFs is critical for motion segmentation. Our finding that perceptual integration of multiple motion directions occurs when spatial separation is less than V1 RF size is also new.

### Key factors that determine motion segmentation

In our visual stimuli, we used a fixed speed (5°/s). As we varied the path length of paired dots, the lifetime of the dots (the duration they were moving before being replotted) was also changed. We did not equate the lifetime by changing motion speed, because speed can influence motion segmentation, and faster speed makes segmentation harder (Masson et al., 1999; Rocchi et al., 2018; Huang et al., 2024). It is known that the motion duration of random dots impacts motion discrimination, integration, and segmentation (e.g., Watamaniuk and Sekuler, 1992; Watamaniuk, 1993; Masson et al., 1999; Bischof et al., 1999; Borghuis et al., 2019). We think that the transition from subjects’ report of “no motion” to “a single motion direction” is likely due to the increase in the path length and dot lifetime. However, we think spatial separation at the scale of V1 RFs is a key factor for the transition from direction integration to segmentation in our experiments. Data in Figure 2 support this notion. For example, as illustrated by the vertical dashed line at the spatial separation of 0.43°, the path length (0.6°) and the dot lifetime (120 ms) were fixed. However, direction segmentation performance progressively decreased as eccentricity increased, from very good at small eccentricity (3°) to poor at larger eccentricity (7°) (Fig. 2). This result can be explained by V1 neurons’ RF sizes at these eccentricities (RF size = 0.35° at 3° eccentricity, and RF size = 0.53° at 7° eccentricity) (based on Dow et al., 1981), relative to the fixed spatial separation of 0.43°. It is known that segregation of transparent motion is worse at peripheral than at central vision (De Bruyn, 1997; Mareschal et al., 2008). However, there is no clear evidence that a longer dot lifetime can rescue segmentation at larger eccentricities when spatial separation is fixed. In fact, the previous finding that multiple motions tend to blend in the periphery can be explained by the spatial separation between motion components not being large enough relative to the V1 RF sizes at those eccentricities.

Additional lines of evidence from previous studies also support the idea that spatial separation, rather than dot lifetime, is the key factor in motion segmentation. In an additional experiment in the study by Qian et al. (1994), they used a fixed path length (0.2°) and a dot lifetime (100 ms) for paired dots that moved horizontally in opposite directions. They found that increasing the vertical spatial separation between the paired dots made the stimulus directions more transparent. In the study by Mestre et al. (2001), they used paired dots that always moved for 200 ms, and found that increasing the spatial separation between the paired dots enhanced speed segmentation.

### Implications for the neural mechanism underlying motion segmentation

This study reveals that spatial separation between motion components at the scale of V1 RFs is essential for motion segmentation, and provides new insights into the neural mechanisms underlying motion segmentation and transparency. A parsimonious interpretation is that when the spatial separation between dots moving in different directions exceeds the RF size of V1 neurons, dots moving in individual component directions would fall into the RFs of different V1 neurons. Therefore, V1 neurons can represent individual component directions, enabling motion segmentation. In contrast, when the maximal spatial separation between paired dots is smaller than the RFs of V1 neurons, the locally paired dots would likely be confined in the RFs of individual V1 neurons. We hypothesize that V1 neurons integrate the component directions of locally paired dots within their RFs into the single VA direction; because this local integration abolishes the signals of individual component directions at the V1 level, it becomes impossible to selectively integrate the local component directions over a long range to achieve motion segmentation and transparency; instead, long-range integration of the VA direction represented by individual V1 neurons would give rise to the perception of an integrated single direction.

Our interpretation and hypothesis imply that V1 plays a crucial role in motion transparency and segmentation, and predict that V1 neurons should represent the VA direction of locally paired dots with spatial separation less than V1 RF size, and represent the individual component directions when the spatial separation exceeds V1 RF size. Preliminary neurophysiology results from our lab are consistent with these predictions (Ghimire et al., 2025 SFN abstract). Our interpretation aligns with previous psychophysical studies on motion segmentation based on speed cues (Masson et al., 1999; Mestre et al., 2001). Masson et al. (1999) showed that speed tuning in perceptual segmentation closely resembles that of V1 neurons, but not MT neurons. Mestre et al. (2001) established that the spatial scale important for speed segmentation matches RF sizes of V1 neurons. Both studies indicate V1’s role in motion segmentation.

Our interpretation, however, differs from the suggestion by Qian and colleagues (Qian et al., 1994a; Qian and Andersen, 1994; Qian et al., 1994b). In a series of elegant studies, Qian et al. were the first to use locally paired dot stimuli to study motion transparency and segmentation. Qian and Andersen (1994) recorded from V1 and MT neurons of macaque monkeys in response to locally paired dots and unpaired dots moving in opposite directions. Unpaired dot stimuli elicit the perception of motion transparency, whereas locally paired dots abolish it. They found that, while MT neurons showed a significant change in the firing rates to the paired and unpaired dots, the responses of V1 neurons did not change significantly. They suggested that the lack of motion transparency with the locally paired dots was due to a local suppression mechanism occurring at the level of MT, and V1 does not play a crucial role in motion transparency (Qian and Andersen, 1994; Qian and Andersen, 1995). We think the discrepancy in the interpretations is related to the motion directions used. When locally paired dots move in opposite directions, perception of flickering noise, rather than motion direction, is elicited (Qian et al., 1994a). The difference in how MT and V1 neurons respond to unpaired and locally paired dots may depend in part on how neurons in those areas respond to flickering noise, rather than reflect whether neurons can or cannot represent the component directions. In the current study, we used two motion directions separated by 90°. Locally pairing dots of these stimuli changed the perception from motion transparency to an integrated VA direction (Curran and Braddick, 2000). Our finding that the spatial scale of this perceptual transition closely matches V1 RF sizes across retinal eccentricities suggests that V1 plays a key role in motion transparency and segmentation.

## Materials and Methods

### Participants

Four subjects (coded initials as RH, BA, AS, SC) participated in this study. AS was a naive subject. The naïve subject was familiar with visual psychophysics tasks but unaware of the purpose of this experiment. All observers had normal or corrected-to-normal visual acuity. All aspects of this study were in accordance with the principles of the Declaration of Helsinki and approved by the Institutional Review Board at the University of Wisconsin-Madison. Subjects provided written informed consent before participating in the experiments.

### Visual display and experimental procedure

All visual stimuli were presented on a 23-inch Planar SA2311W LED monitor, which operated at a 100 Hz refresh rate to ensure smooth motion perception and a spatial resolution of 1920 × 1080 pixels for precise stimulus rendering. Participants were seated at a fixed viewing distance of 60 cm from the monitor. At this distance, the screen subtended a visual angle of approximately 49.3° × 29.5°, and each pixel subtended approximately 0.029°, yielding a display resolution of about 34.5 pixels per degree of visual angle. The experiment was controlled by custom scripts written in MATLAB (R2015a; MathWorks, Natick, MA), utilizing the Psychophysics Toolbox 3 extensions (Brainard, 1997; Pelli, 1997). This software suite is standard in vision research and allowed for the precise timing and contrast control necessary for rendering the dynamic stimuli according to the specified parameters.

The experiment took place in a dark room with dim, indirect background illumination. To ensure a stable head position and a constant viewing distance throughout the experiment, participants were positioned in a chin rest with a forehead support. This stabilization was critical for maintaining the accuracy of the retinal eccentricity manipulation, as any head movement would alter the location of the stimulus on the retina. The monitor’s luminance output was linearized through gamma correction. We used a Minolta LS-110 photometer to measure the luminance for a range of grayscale values and generated a lookup table to ensure that the relationship between the requested pixel intensity in the software and the actual light output from the monitor was linear. For this experiment, the background was set to a uniform mid-gray (40 cd/m2), and the white dots had a luminance of 80 cd/m2. A small red fixation dot (0.2° diameter) was presented at the center of the screen.

To ensure that participants maintained stable fixation on the central fixation dot, thereby ensuring the stimuli were presented at the intended retinal eccentricities, we continuously monitored eye position using a Tobii Pro Nano Eye Tracker (Tobii AB, Sweden). The device, which sampled gaze position at 60 Hz, was mounted to the bottom of the monitor. Before each block of trials, a five-point calibration procedure was performed using the Tobii Eye Tracker Manager software to map the participant’s gaze to the screen coordinates accurately. During each trial, eye tracking data was analyzed in real-time. A circular tolerance window with a radius of 0.5° of visual angle was established around the fixation dot. If the participant’s gaze deviated outside this window for more than 50ms during the stimulus presentation, the trial was immediately aborted and recycled to be presented again later in the block. This strict fixation-contingent paradigm ensured that all analyzed data came from trials where the stimulus eccentricity was properly controlled.

### Visual stimuli

The paired-dot stimuli are composed of locally paired dots that move in two different directions with a limited lifetime (Qian et al. 1994a; Curran & Braddick, 2000) (Fig. 1A). Two locally paired dots were rendered such that the stimuli first appear in the aperture with a fixed maximum spatial separation, denoted as λ, and move in two directions at the same speed v, initially in a converging course. After the two dots intersect, they continue to move in a diverging course until they complete the total pathlength. The dot pair was then be turned off and reappeared in a different location within the aperture until the end of the motion period. The maximum spatial separation λ = ρ• sin(α/2), where α is the direction separation (DS) between two motion directions (DS = 90º) and ρ is the corresponding pathlength of each dot. We created stimuli with varied path length (ρ) traversed by each dot, and hence varying the maximal spatial separation (λ) between the dots in each pair.

### Experimental design and procedure

We manipulated two stimulus parameters. First, we varied the path length (ρ) traversed by each dot, and hence the maximal spatial separation (λ) between the dots in each pair. The path length was varied from 0.1° to 1.0° in steps of 0.1°, corresponding to maximal spatial separations of 0.071° to 0.707°. Second, we centered the stimulus aperture at five different retinal eccentricities ranging from 3° to 7° (3°, 4°, 5°, 6°, or 7°) from the fovea (Fig. 1B) in separate, randomized sessions. Within each session, trials with different spatial separations were presented in a random order. Each trial began with the presentation of the central red fixation dot. After a 300 ms static presentation of the dot field, the dots moved for 700 ms. Following the motion period, the screen went blank, and participants had 2000 ms to report their percept by pressing one of three designated arrow keys. Subjects performed a three-alternative forced-choice (3AFC) task, reporting whether the visual stimulus had no motion, moved in a single direction, or moved in two directions (Fig. 1C). Each participant completed 6-11 sessions for each eccentricity.

### Data analysis

#### Psychometric function fitting

To derive the critical spatial separation (λ_75_) for each experimental session, we fit a logistic function to the proportion of trials in which the observer reported perceiving two directions as a function of spatial separation. The logistic function was defined as:

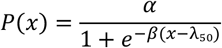

where *P*(*x*) is the proportion of two directions responses, *x* is the spatial separation, *α* is the upper asymptote (performance ceiling), *β* represents the slope, and λ_50_ is the threshold at the inflection point (50% of the asymptote). Fits were performed using a least-squares procedure (MATLAB *lsqcurvefit* function). Goodness-of-fit was assessed using the coefficient of determination (R^2^). As all sessions yielded reliable fits, no session data were excluded from the analysis.

#### Critical spatial separation (λ_75_)

We defined the critical maximum spatial separation (λ_75_) as the spatial separation corresponding to 75% of the performance (asymptote). This metric was chosen to represent a robust threshold for reliable perception of two directions, well above the point of subjective equality. λ_75_ was derived from the fitted parameters λ_50_ and *β* using the inverse logistic function:

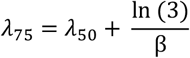

These derived λ_75_ values served as the input for all subsequent statistical analyses.

#### Linear mixed-effects modeling

To quantify the effect of eccentricity on segmentation thresholds while explicitly accounting for the hierarchical structure of the data, we analyzed session-level threshold estimates using a Linear Mixed-Effects (LME) model. This modeling approach allowed us to decompose the variance in performance into three distinct components: Eccentricity (Fixed Effect): The primary experimental manipulation. Participants (Random Effect): To account for inter-subject variability, we included random intercepts and random slopes for each participant. Sessions (Residual Error): By analyzing individual session estimates rather than subject averages, the model preserved session-to-session fluctuations as residual variance.

The model was implemented in MATLAB using the *fitlme* function (Statistics and Machine Learning Toolbox) with the following specification:

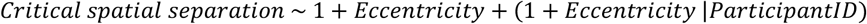

This corresponds to the formal equation:

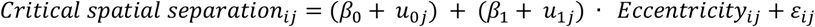

Where *Critical spatial separation*_*ij*_ is the λ_75_ estimate for the *i* − *th* session of the *j* − *th participant* ; β_0_ and β_1_ are the fixed population intercept and slope; *u*_0*j*_ and *u*_1*j*_ are the random intercept and slope deviations for participant *j*; and ε_*ij*_ is the residual error for the specific session. The model’s explanatory power was quantified using Adjusted *R*^*2*^.

#### Bootstrapping and descriptive statistics

To robustly estimate the precision of critial spatial separation measurements for data visualization (Fig. 3A), we employed a non-parametric bootstrap procedure. For each experimental condition (participant × eccentricity), the collected session-level critical spatial separations (λ_75_) were resampled with replacement 1,000 times to generate a bootstrap distribution. The arithmetic mean of the original data served as the point estimate. Symmetric error bars representing the 95% confidence interval (CI) were calculated as ±1.96 × Bootstrap_SD, where Bootstrap_SD is the standard deviation of the bootstrap distribution. This method avoids assumptions about the normality of session-to-session variability. Validation confirmed that bootstrap means converged closely with arithmetic means (mean absolute difference ∼ 0.002°).

